# Machine-assisted cultivation and analysis of biofilms

**DOI:** 10.1101/210583

**Authors:** Silla H. Hansen, Tobias Kabbeck, Carsten P. Radtke, Susanne Krause, Eva Krolitzki, Theo Peschke, Jannis Gasmi, Kersten S. Rabe, Michael Wagner, Harald Horn, Jürgen Hubbuch, Johannes Gescher, Christof M. Niemeyer

**Affiliations:** Institute for Biological Interfaces (IBG-1), Karlsruhe Institute of Technology (KIT), Herrmann-von-Helmholtz Platz 1, D-76344 Eggenstein-Leopoldshafen, Germany.; Institute for Applied Biosciences; Institute of Engineering in Life Sciences, Section IV: Biomolecular Separation Engineering (BLT-MAB), Fritz-Haber-Weg 2, D-76131 Karlsruhe.; Engler-Bunte-Institute for Water Chemistry and Water Technology, Engler-Bunte-Ring 5, D-76131 Karlsruhe, Germany.

**Author notes:** Correspondence and requests for materials should be addressed to J.G. or C.N.

## Abstract

Biofilms are the natural form of life of the majority of microorganisms. These multispecies consortia are intensively studied not only for their effects on health and environment but also because they have an enormous potential as tools for biotechnological processes. Further exploration and exploitation of these complex systems will benefit from technical solutions that enable integrated, machine-assisted cultivation and analysis. We here introduce a microfluidic platform, where readily available microfluidic chips are connected by automated liquid handling with analysis instrumentation, such as fluorescence detection, microscopy, chromatography and optical coherence tomography. The system is operable under oxic and anoxic conditions, allowing for different gases as feeding sources and offers high spatiotemporal resolution in the analysis of metabolites and biofilm composition. We demonstrate the platform’s performance by monitoring the self-organized separation of mixed cultures along autonomously created gradients, the productivity of biofilms along the microfluidic channel and the enrichment of microbial nanoorganisms.

## Introduction

Spurred by the need of fundamental and applied research in chemistry and biomedicine to more efficiently use human resources and to improve safety, efficacy, and scope of development and production processes, the implementation of machine-assisted approaches is currently attracting great attention. The automated conduction of solid-phase reactions aided by fluid handling used therein has already led to impressive advances in chemical synthesis programms^1–4^ and the manufacturing of pharmaceutical compounds^5, 6^. These developments are complemented by current advances in the design of microfluidic lab-on-a-chip systems for applications in the life sciences, which include fundamental studies of cellular processes, biomedical diagnostics or drug discovery^7–10^. Biofilms are the natural form of life of the majority of microorganisms. These multispecies consortia are intensively studied for their effects on health and environment and they also hold potential as tools for biotechnological processes^11–13^. Several studies have applied microfluidic techniques to the field of biofilm analysis, primarily to investigate biofilm formation of model organisms that may even be genetically programmed^14–19^. However, applied and basic biofilm research is redirecting its focus away from tractable model systems and will depend on robust, automated solutions for multivariate analysis of mixed species biofilms containing potential novel and superior biocatalysts for biotechnological processes^13^. As a step towards this goal, we here report on the development of an integrated platform for machine-assisted biofilm research. The system is based on microfluidic PDMS chips mounted on tailored interfaces to connect with hardware for automated liquid handling and instrumental analysis, such as fluorescence reading, epifluorescence microscopy, optical coherence tomography (OCT) and liquid chromatography (Figure 1). The platform is operable under oxic and anoxic conditions, allowing for different gases as feeding sources and offers high spatio-temporal resolution in the analysis of metabolites and biofilm composition. We demonstrate the platform’s applicability and versatility by investigation of the self-organized separation of mixed cultures along autonomously created gradients, biofilm-catalyzed stereoselective transformations, and the enrichment of acidophilic nanoorganisms. We believe that the here presented implementation of machine-assisted microfluidics, robotic handling, and in-depth instrumental analysis is an important advance for the exploration and exploitation of biofilm development and community dynamics.

**Figure 1:**
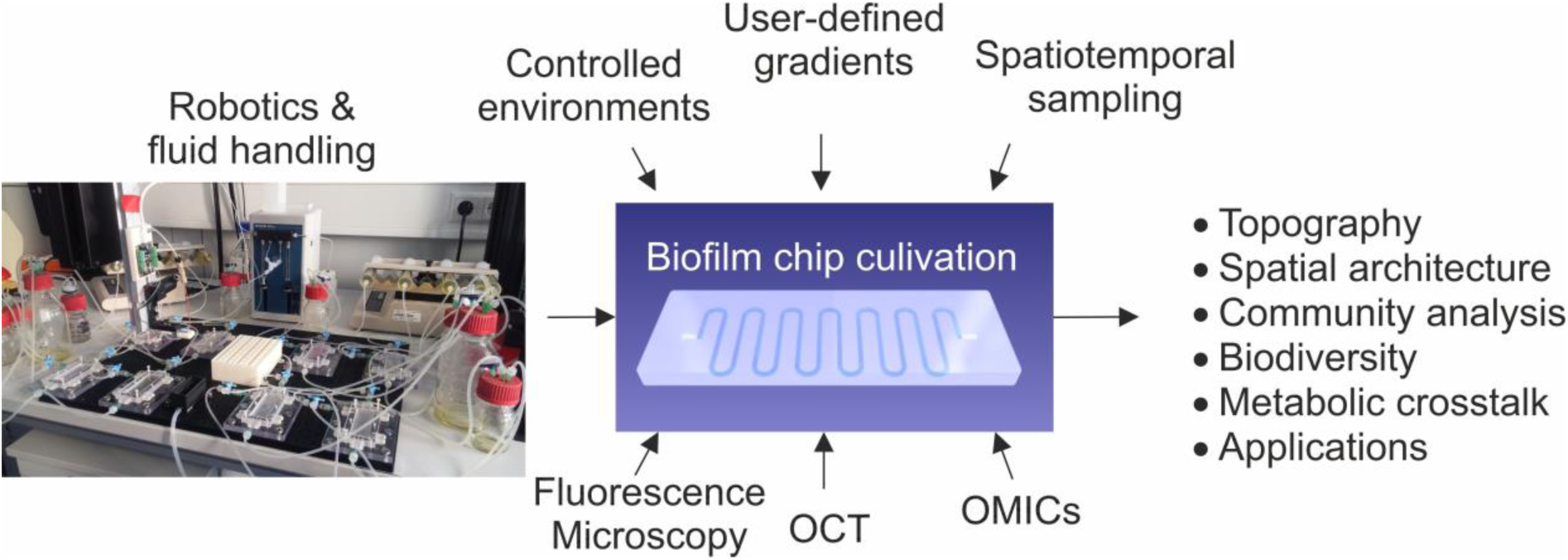
Overview of the integrated platform for machine-assisted cultivation and analysis of biofilms.

## Results

### The automated platform for biofilm cultivation and analysis

As illustrated in Figure 1, our platform is based on microfluidic channel structures, which can be readily produced from the elastomer polydimethylsiloxane (PDMS) by replica casting using micromolds prepared by micromachining or photolithographic techniques^20^. The PDMS chips are sealed with coverslips to enable the accurate microscopic inspection of the microchannel’s interior. The fluidic chips are mounted in cartridges, which serve as the interface to pumps and valves for media supply to enable cultivation of biofilms under controlled conditions for prolonged periods; so far, we conducted cultivation campaigns of up to 4 months. Non-destructive *in situ* imaging of living biofilms was performed by means of light microscopy (incl. epifluorescence) and optical coherence tomography (OCT)^21^. Since PDMS is gas permeable, a casket was developed, which allows long-term anoxic cultivation as well as the use of arbitrary gas mixtures as carbon and electron sources (Supplementary Figure S1). Furthermore, to enable online measurement of oxygen inside the microfluidic flow cells, methods for the integration of fiber-optical sensors were developed (Supplementary Figures S1, S2). The cartridge also functions as the interface to a liquid handling station (LHS, Supplementary Figures S3, S4), a widely established automation tool in biological laboratories. Here, the LHS was utilized to enable automated end-point analysis of the biofilms by fluorescence *in situ* hybridization (FISH) or catalyzed reporter deposition FISH (CARD-FISH), as discussed below. Importantly, the standardized technical interface allows to follow the automated FISH procedures by imaging techniques like OCT, for instance, to quantify mechanical abrasion during the various washing and incubation steps of the FISH procedure (Figure S5).

A central instrument of the developed platform is a robotic sampler that allows to repeatedly draw small liquid sample volumes (1-10 µL) directly from arbitrary positions of the microfluidic channel without interruption of a running cultivation (Figure 2, see also Supplementary Figures S6-S10). The sampler is equipped with a sharp cannula, connected to pumps for sample extraction, which is freely movable in either X-, Y- or Z-direction (600 mm × 300 mm × 200 mm, respectively) with a precision of ±25 μm in X- and ±10 μm in Y- and Z-direction. Exact positioning of the cannula is controlled by an automated pattern recognition software (Supplementary Figure S7) to assure precise repeated sample drawing over the entire experiment. The vertical positioning of the needle is controlled by a pressure sensor (Supplementary Figure S8) to enable the puncturing of the PDMS layer on top of the channel, withdrawal of either liquid from above or cell material from within the biofilm, and subsequent delivery of the samples to microplates. After sampling, the flexible PDMS layer seals back to leave a closed channel that enables continued cultivation. The collected samples are transferred to a cooled storage compartment (Figure S9) from which they can be subjected to further analysis by e.g., chromatography or sequencing (discussed below). The robotic sampler is controlled by a graphical user interface that uses prescribed script modules to conduct the various procedures necessary for automated sample extraction from the microfluidic chips (Figure S10).

**Figure 2:**
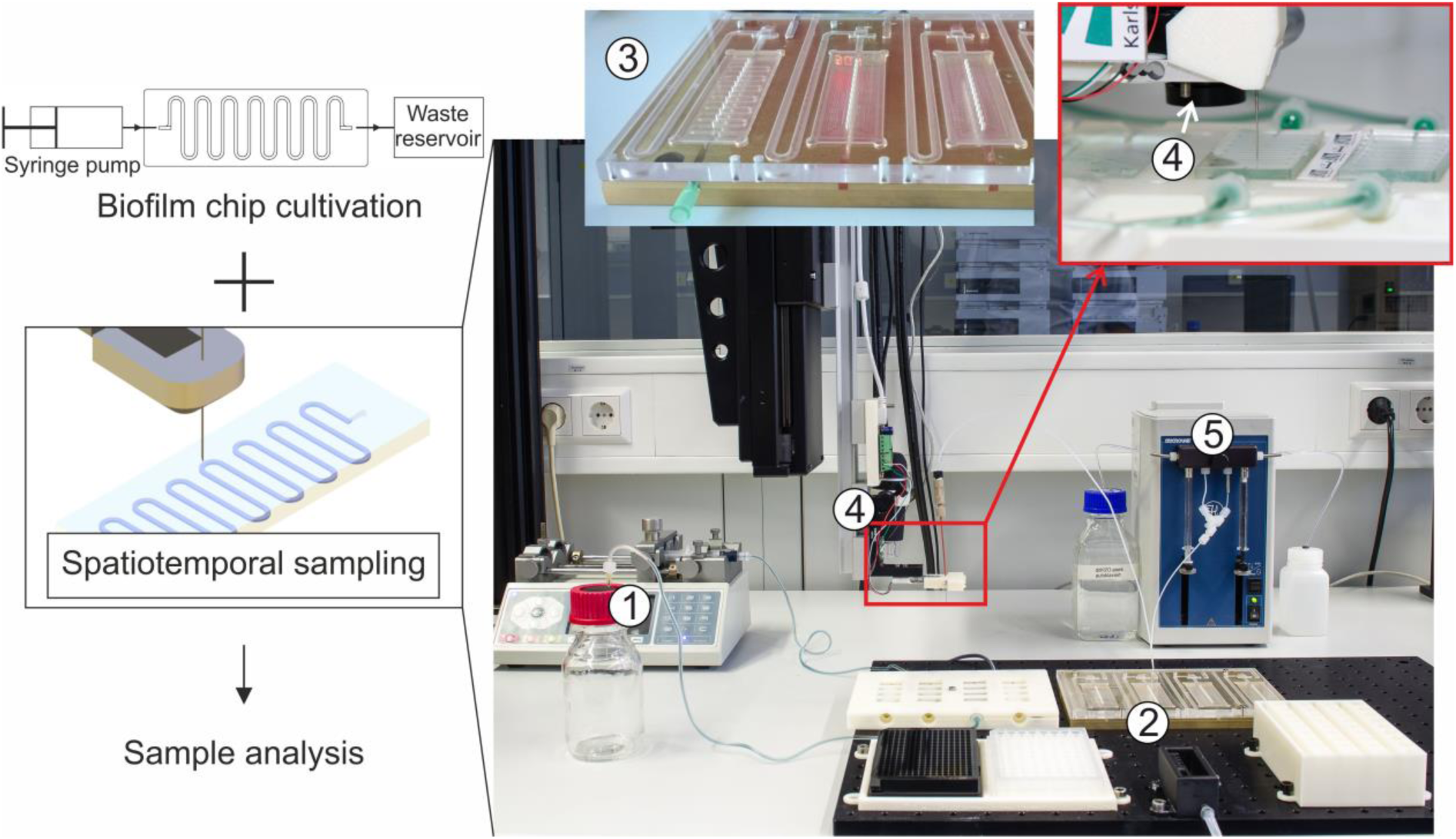
Robotic sampler for non-destructive spatiotemporal *in situ* analysis of flow cell-cultivated biofilms. The overview image shows external syringe pumps (1) used for continuous perfusion of the microfluidic bioreactors with medium or substrate, the robotic deck of the sampling device (2) onto which a custom-made temperature-controlled chip holder is mounted (3), the sampling head (4) that is connected to the pumping unit (5) for withdrawal of small sample volumes from the channel. Further details on the design of hardware parts and control software are shown in Supplementary Figures S6-S10.

### Implementation of automated FISH procedures for endpoint analysis of flowcell cultivated biofilms

FISH or CARD-FISH are the gold standard in biofilm research because they allow for the analysis of the composition of microbial communities with an extraordinarily high spatial resolution, multiplexing capability and sensitivity^22, 23^. A disadvantage of this technique is that the experimental execution of FISH requires multiple, time consuming steps for labeling of the ribosomal RNA (rRNA) targets with fluorescent oligonucleotide probes. To overcome this obstacle and motivated by approaches to the semi-automated conduction of FISH for eukaryotic cell culture^24–26^, we developed a technical interfaces that enabled implementation of automated FISH/CARD-FISH analyses into our robotic platform. Based on a previous design^27^, the dimensions, contact points and connections of our flowcell-to-LHS interface (Supplementary Figure S4) are compatible with standard microtiter plate equipment, such as thermocyclers, shakers and plate readers. Hence, the automated system conducts all steps of the FISH procedure autonomously. To demonstrate the system’s utility, we carried out fully automated FISH analyses of single- and mixed-species biofilms of the gram-negative organism *Escherichia coli* and the gram-positive strain *Bacillus subtilis* that were grown in nine independent linear flow chips with constant medium supply (Figure 3).

**Figure 3:**
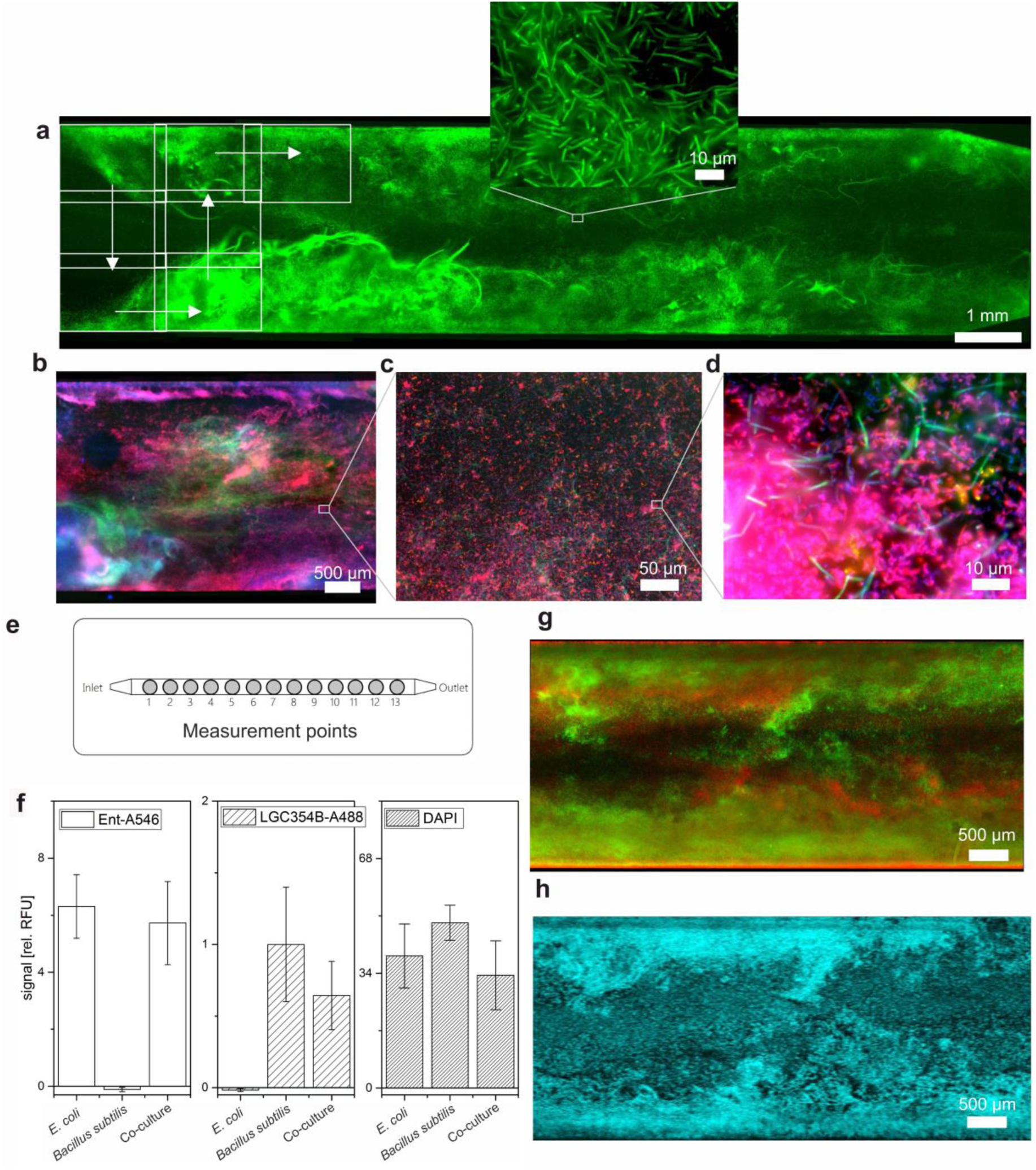
Optical analysis of flowcell biofilms labeled by automated FISH. Overview image and high-resolution (inset) micrographs of a pure *B. subtilis* (a) and a mixed species *E. coli* and *B. subtilis* biofilm of (b-d). Biofilms were grown in triplicates in linear flow chambers (e), subjected to automated FISH and analyzed with the integrated fluorescence reader at the 13 measurement points to obtain mean fluorescence values for the nine parallelly cultivated biofilms (f). *B. subtilis* and *E. coli* were labeled with probes LGC354B-A488 (colored in green) and Ent-A546 (red), respectively. Note that DAPI staining indicates approximately equal growth density and FISH data allow for reliable identification of pure *E. coli* and *B. subtilis* cultures. (g) Epifluorescence micrographs of mixed species *B. subtilis* and *E. coli* biofilms (same color code as in a-d) confirm the topographical biofilm features visualized by means of OCT (h). Displayed height = 100 μm (cyan colored).

With this setup, it was possible to characterize the biofilms with the LHS-integrated fluorescence reader (see Supplementary Figure S3) as well as to image them by epifluorescence microscopy. The analysis of predefined spots inside the linear channels with the fluorescence reader allowed to clearly distinguish between single and mixed species biofilms (Figure 3 e/f). However, owing to the limited lateral resolution of conventional plate readers, the accurate and quantitative mapping of biofilm composition required high-resolution fluorescence imaging. To this end, sequential images were taken (indicated by rectangles, in Figure 3 a) and merged afterwards by grid stitching^28^ to document with high resolution large areas of up 3 × 14 mm inside the flowcell. The method allows to reveal mesoscopic features, such as the streamers formed by the *B. subtilis* inside the broader context of the flowcell (Figure 3 a), as well as the microscopic composition of mixed species biofilms (Figure 3 b-d), thereby clearly illustrating the advantages of the developed method for multiscale FISH imaging of native biofilms directly inside their cultivation vessel. Furthermore, comparison of OCT images with the final FISH images revealed that biofilm features visualized by OCT are easily recognizable by epifluorescence microscopy after FISH (Figures 3 g, h). The implementation of OCT also clarified that only slight biofilm detachment occurred during the automated FISH procedure but the overall structural integrity was preserved (Figure S5).

### Enrichment of rare microbial species

We investigated the performance of our platform for selective enrichment of rare species in natural biofilms, such as ARMAN (Archaeal Richmond Mine Acidophilic Nanoorganisms). The inoculum was an acid mine drainage (AMD) biofilm (Figure 4 a) composed of acidophilic autotrophic bacteria that inhabit the outer oxic rim of the biofilm and archaea that thrive in the anoxic center^29^. Oxic medium was perfused through the chip that was mounted inside the developed gasket (Figure S1 h) and flushed with a mixture of N_2_ 80% /CO_2_ 20%. We could measure a depletion of oxygen in the flow channel that led to anoxic medium conditions at the outflow. The slow growth of the organisms demanded long cultivation times of 4 months. Imaging of the cells by CARD-FISH revealed that the growth conditions led to a selection for archaea that were underrepresented in the inoculum but dominated growth within the chip (Figure 4 b). However, we could not show growth of ARMAN using whole AMD biofilm samples as inoculum.

**Figure 4:**
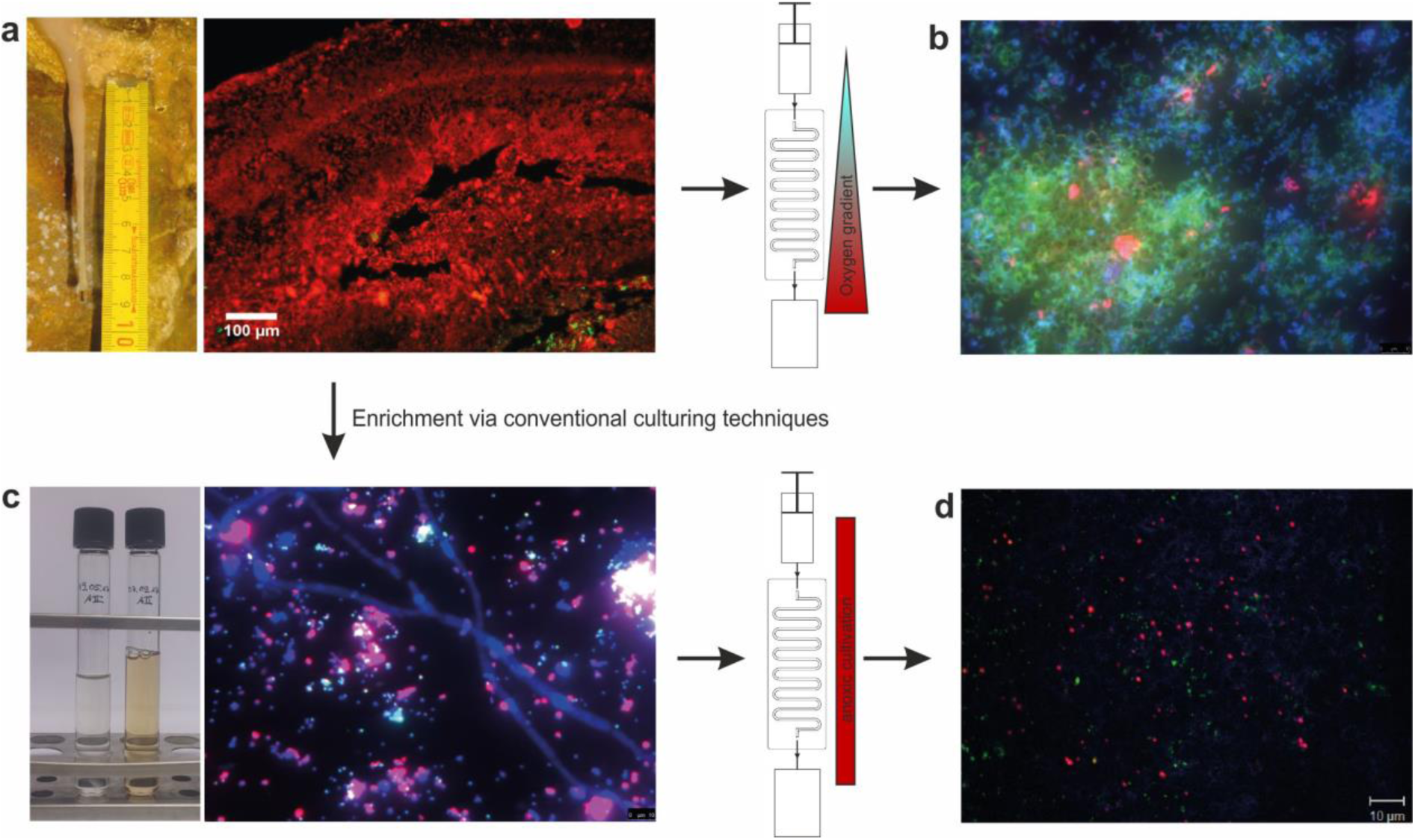
Enrichment of acidophilic archaea. (a) Native AMD-biofilm and CARD-FISH picture of a cross section. (b) CARD-FISH picture of highly enriched archaea grown in the microfluidic channel that was inoculated with the native biofilm and cultured in an oxygen gradient. Bacteria stained in red (EUB-I, Alexa546), archaea in green (Arch915, Alexa488) and DAPI stained cells in blue. (c) CARD-FISH image of an enrichment culture containing acidophilic Archaea, ARMAN-related organisms and a fungus. (d) CARD-FISH picture of an enrichment culture grown in the microfluidic device, containing only acidophilic archaea and ARMAN-related organisms. Archaea stained in red (ARCH915, Alexa546), ARMAN in green (ARM980, Alexa488) and DAPI stained cells in blue. Note that ARMAN, although belonging to the Archaea, cannot be detected with the general ARCH915 probe.

Therefore, fluidic cultivation of a stable laboratory culture of anaerobic acidophilic nanoorganisms^30^ was conducted. This ARMAN co-culture was derived from the aforementioned acidophilic environmental AMD biofilm (shown in Figure 4 a) and evolution by several transfers in anoxic medium under planktonic conditions led to a community that was composed of three novel archaea including an ARMAN species and a fungus^30^ (Figure 4 c). We used this co-culture as inoculum for microfluidic cultivation inside the anoxic gasket. Using CARD-FISH and 16S-rRNA analysis, we found that fluidic cultivation was successful. Indeed this technique enabled the cultivation of the first co-culture that contained only an ARMAN species along with a single helper organism (Figure 4 d), dubbed as B_DKE that we had previously observed in the more complex co-culture obtained by conventional cultivation techniques^30^. Most likely, B_DKE is necessary as producer for compounds that cannot be produced by the multi-auxotrophic ARMAN organism. We hypothesize that the further enrichment obtained with the microfluidic platform is attributable to the implementation of biofilm conditions for culturing, which comprise the natural form of growth of these organisms. Systematic variation of culturing conditions should now open the door to the in-depth analysis of distinctive interspecies dependencies.

### Mutually dependent two-species biofilms

As a result of microbial activity and/or the diffusion of gases out of or into the medium via the PDMS materials, the microfluidic channels of our platform allow for the formation of gradients along the medium flow. This feature can be exploited for the cultivation of mutually dependent communities that are fluidically connected but spatially separated. The self-organized separation of mixed cultures along autonomously created gradients can be controlled and steered by the applied growth conditions and it may also allow to determine specific characteristics of novel environmental isolates, such as growth auxotrophies that are difficult if not impossible to study by conventional approaches.

As a proof of concept, we studied the interaction of a chromate resistant *Leucobacter chromiiresistens* strain with the laboratory model organism *E. coli*. *L. chromiiresistens* has the ability to tolerate the highest chromate concentrations reported so far^31, 32^. It is an aerobic organism whose chromate tolerance stems from its capability to efficiently reduce toxic Cr(VI) to Cr(III) species, the latter of which are highly insoluble and therefore less toxic. We hypothesized that flow cultivation of mixed *E. coli*/*L. chromiiresistens* cultures with Cr(VI)-containing medium should lead to spatial enrichment of *E. coli* in the downward sections of the chip, while the channel’s front section should be predominantly inhabited by *L. chromiiresistens*. To achieve an optimal characterization of the biofilm composition, we took advantage of both the automated FISH procedure and the robotic sampler. The latter was used to draw 10 μl samples from the biofilms, which were subjected directly to 16S rRNA gene amplification and subsequent Ilumina sequencing.

We conducted parallel co-cultivation experiments in LB-medium that contained variable chromate concentrations ranging from 0-3 mM. In the absence of chromate, the biofilm consisted almost exclusively of *E. coli* cells throughout the entire flow channel (Figure 5 a). Hence, in the absence of toxic chromate, the higher growth rates of *E. coli* led to overgrowth of *L. chromiiresistens* species. In the presence of 1 mM chromate, formation of two separated growth areas was observed with an abundance of *L. chromiiresistens* and *E. coli* species at the front and rear regions of the channel, respectively (Figure 5B). While *L. chromiiresistens* dominated the oxic part containing the highest chromate concentration, *E. coli* could thrive in the rear parts of the channel presumably due to the decrease in Cr(VI) concentration and its ability to thrive as facultative anaerobic organism. A further increase of the chromate concentration led to biofilms that were dominated by *L. chromiiresistens* cells throughout the channel (Figure 5 c, d). However, in the case of 2 mM Cr(VI) concentration, 16S amplicon sequencing still clearly revealed preferred *E. coli* growth at the rear end of the channels. These data indicate that the amount of *L. chromiiresistens* biomass grown in the channels’ front part has a sufficient capacity to reduce about 1 mM Cr(VI) while, at higher concentrations, the remaining chromate could be no longer efficiently removed to allow growth of *E. coli*. Hence, the results provide a clear demonstration that mutually dependent consortia self-organize along an autonomously created chemical gradient into spatially separated populations. Furthermore, preliminary experiments carried out with aerobic fumarate reduction deficient *E. coli* Δfrd and anaerobic fumarate reducing *G. sulfurreducens* biofilms confirmed the self-organization of mutually dependent consortia along the growth-dependent chemical gradients of oxygen and organic acids created by *E. coli* (Supplementary Figure S11).

**Figure 5:**
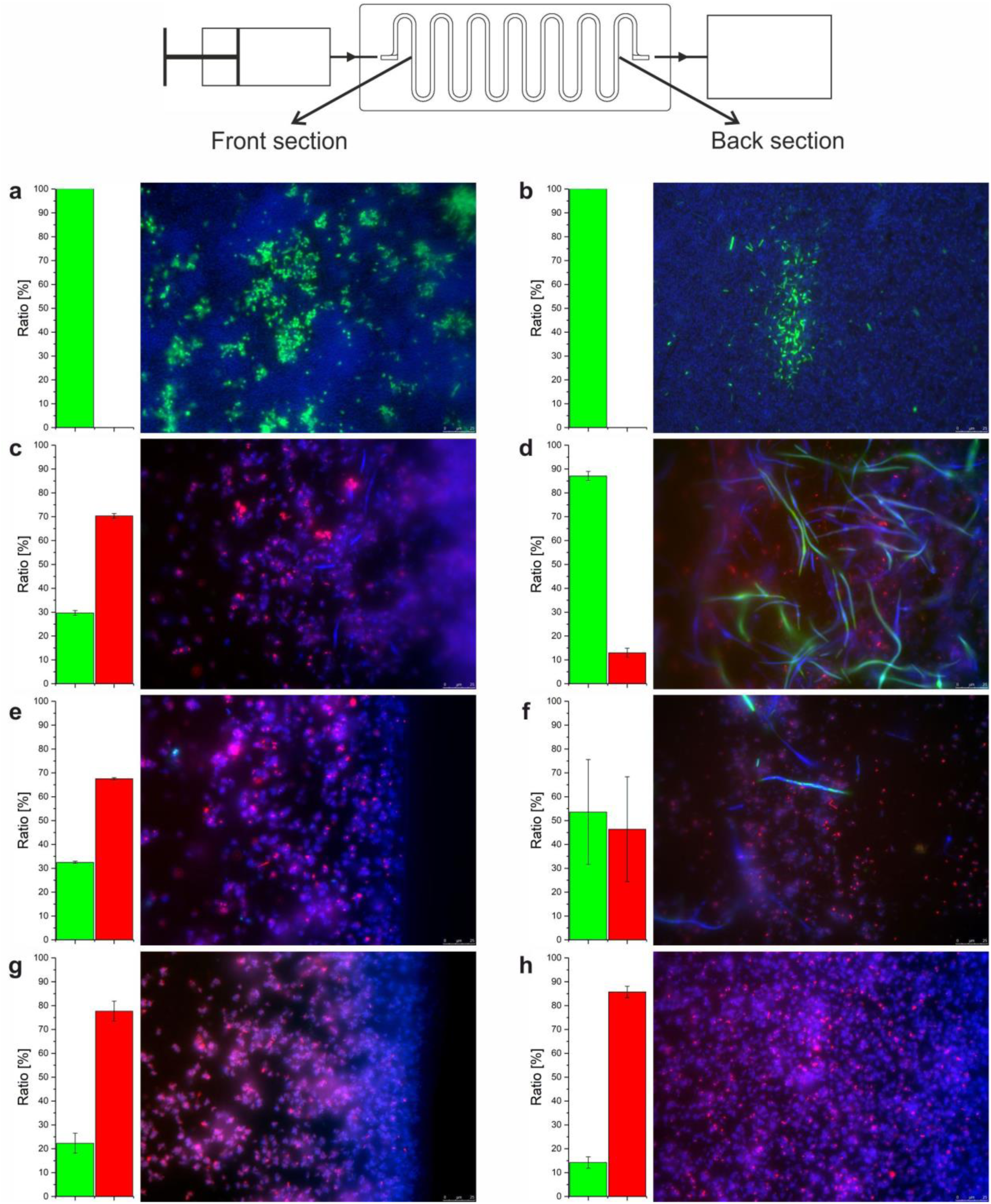
Mutually dependend biofilms consortia self-organise along an autonomously created gradient of chromate. A series of four identical inocula was used and cultivated over 7 days in the absence (a, b), or the presence of 1 mM (c, d), 2 mM (e, f) or 3 mM (g, h) chromate containing medium. Epifluorescent images and phylogenetic composition in (a), (c), (e), (g) represent the front section of the microfluidic chip while (b), (d), (f), (h) show representative data from the rear end. Sampling points are indicated in the scheme of the setup. *Escherichia coli* cells are shown in green, *Leucobacter chromiiresistens* cells in red. Unspecific labeling with DAPI is indicated in blue. The bar charts on the left side of each sample represent the community composition derived from 16S amplicon sequencing.

### Productive biofilms

Applied biofilm research is concerned with the exploitation of bacterial communities containing novel and superior biocatalysts for biotechnological processes^13^. Hence, to illustrate the scope and utility of our platform, we performed microfluidic cultivation experiments with of a productive *E. coli* biofilm that expresses the *R*-selective alcohol dehydrogenase LbADH from Lactobacillus brevis ATCC 14869. As a test case for continuous biofilm cultivation with a concomitant stereoselective transformation reaction, we used the sequential biocatalytic reduction of 5-nitrononane-2,8-dione (NDK, in Figure 6 a) by LbADH that leads to selective production of the hydroxyketones HK1 and HK2, which are then converted into the pseudo-C2-symmetric *R*,*R*-configured diol product^33, 34^. Using the robotic sampler, samples were taken from 12 different points along the meandric cultivation channel. Analysis by chiral HPLC clearly revealed that the NDK educt vanished while the HK intermediates and diol product evolved with increasing path length (Figure 6 b-d). Together with the results from the metabolic analysis of mutually dependent biofilms described above, this data clearly illustrates the suitability of our platform for multivariate analysis of complex biofilms.

**Figure 6:**
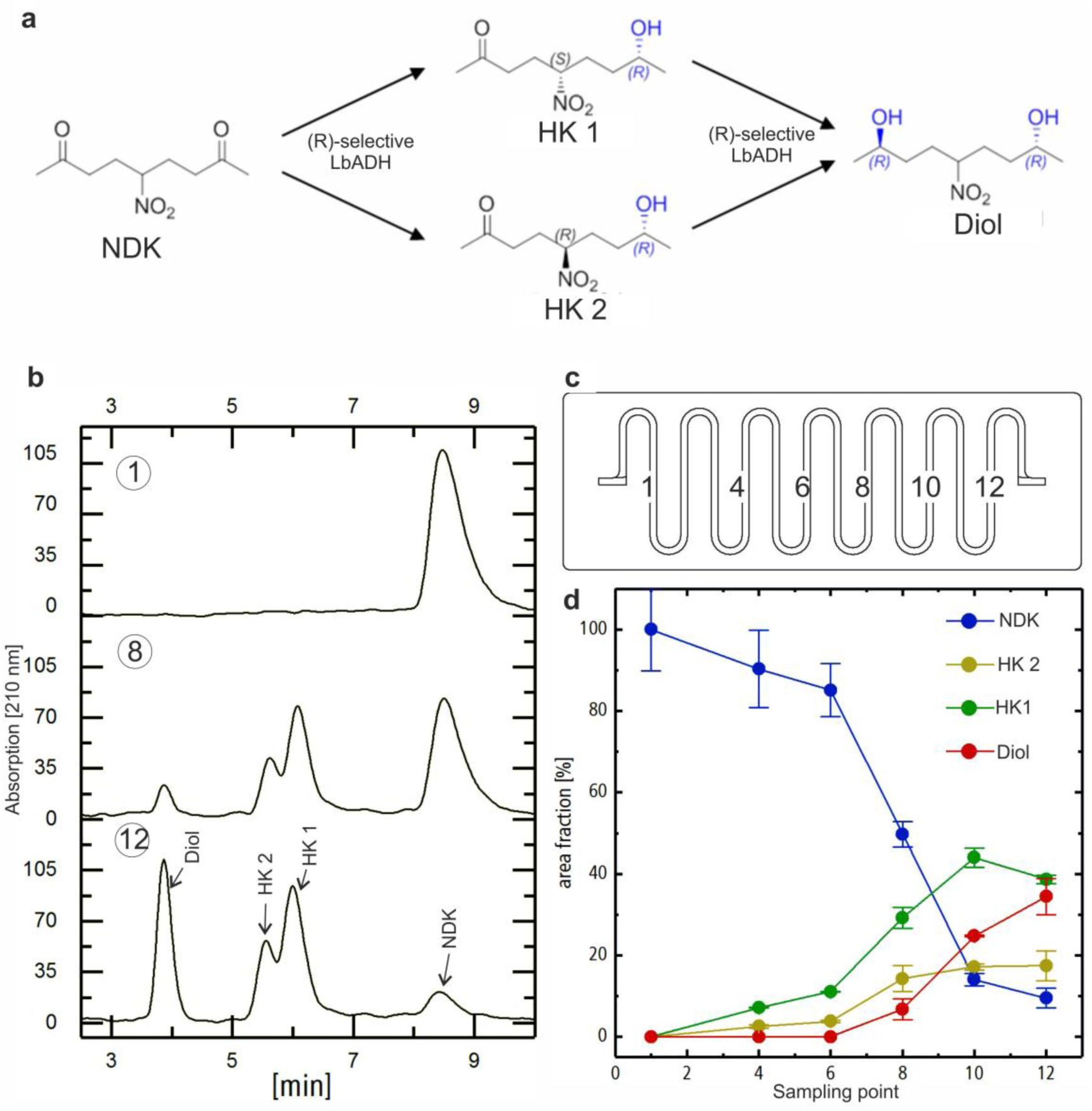
Culturing and analysis of a productive E. coli biofilm. (a) Reaction scheme of the stereoselective reduction of the prochiral nitro-diketone substrate (NDK) into hydroxyketone (HK) and diol products by the *R*-selective ketoreductase LbADH, expressed in a fluidically-cultivated *E. coli* biofilm. (b) Representative chiral HPLC chromatograms of samples drawn from selected points of the meandric cultivation channel (c). The decrease of NDK educt and HK/diol products is clearly evident from the analysis of the reaction samples retrieved along the flowpath at a constant flowrate of 2 µL/min (d).

## Discussion

The here presented platform for machine-assisted cultivation and analysis of biofilms is ideally suited to advance basic and fundamental research on synthetic as well as natural multispecies biofilms. Cultivation can be pursued under arbitrary environmental temperature and gas phase conditions to mimic a large variety of natural habitats. Importantly, our system allows for culturing under flow conditions to establish shear forces that are an essential factor for controlling growth in native environments. The standardized dimensions of the flowcell chips allow for the facile integration of a large variety of designs that are readily accessible by rapid prototyping and soft lithography and can be conveniently adjusted in their length, curvature and internal channel dimensions. This enables diverse applications ranging from rapid growth experiments, as exemplified here with binary *E. coli* and *B. subtilis* cocultures (Figure 3), to long-term studies of the growth of environmental biofilms (Figure 4) and the self-organized separation of mixed cultures along autonomously created gradients (Figure 5). A very important feature of the flow chips is their technical connectivity to commercial liquid handling systems and established analytical instrumentation for imaging. We used a combination of these opportunities to develop automated FISH- and CARD-FISH protocols that unburden the user from time consuming experimental steps and even allow for increased throughput by multiplexing chip-based cultivation experiments. OCT analysis was also implemented in the workflow for online monitoring of biofilm growth and to correlate the fluorescence images to the *in vivo* situation before fixation and labeling. The OCT controlled automation increases the validity of the acquired images as the scientists is no longer a source for operational errors. While automated microscopy can conveniently be implemented for image analysis, it might not even be necessary in certain applications because a plate reader output was shown to reliably help to distinguish between different biofilm-populations.

The opportunity to continuously grow biofilms in the flow cells together with the automated analysis of the biofilm population were exploited to isolate the first binary co-culture of an acidophilic nanoorganism (ARMAN) and a potential helper organism. Previous studies suggested that the reduced genome of ARMAN species will necessitate the activity of other organisms to mask the effect of auxotrophies.^30^ Now, we are able to study the interplay between ARMAN and the helper organism in a highly defined system that can be operated over long periods of time. For this purpose, the advanced highly accurate sampling robot will prove particularly useful. Retrieving samples from defined positions of the cultivation flowcell opens the door for the detailed spatiotemporal analysis of the biofilm. While we used this device and proved its suitability by taking samples for HPLC analysis and amplicon sequencing, one can envision that it can also be employed for implementation of other analytical methods, such as mass spectrometry, or else for the spatiocontrolled injection of substances into living biofilms. This will open the door to study the impact of, for instance, second messengers or antibiotic substances.

16S rRNA gene amplicon sequencing was used to study a mutually dependent synthetic two species biofilm of *Leucobacter chromiiresistens* and *E. coli*. Changing compositions of the biofilm along the flowpath as well as with varying feed medium could be observed. It was also confirmed by subsequent FISH analysis that *Leucobacter* created a niche for *E. coli* cells in the rear part of the flowcell by the reduction of toxic chromate. Furthermore, it could be shown how the online sampling in combination with HPLC can be used to determine concentration levels of metabolic compounds or relevant products. On the one hand, this approach gives access to the composition and dynamics of the microbial community cultivated within the flowcells and, on the other hand, it demonstrates the possibility to evaluate online process parameters for biocatalytic transformation reactions. The latter is considered one of the basic prerequisites for the breakthrough of the use of productive biofilms in biotechnological applications.

In summary, mircofluidic biofilm cultivation devices and tailored robotic instrumentation were combined with powerful standard methods such as (CARD-) FISH, HPLC and next generation sequencing to create a unique set of tools for multispecies biofilm research. Emerging research fields, for example, studies on productive biofilms or the exploration of complex native consortia will largely benefit from the developed platform. It is therefore anticipated that this work represents an important step towards the realization of machine-assisted processes for the next generation of biotechnology.

## Methods

### Chip fabrication

The microfluidic chip designs were based on the dimension of standard microscope slides (76 × 26 mm² DIN ISO 8037-1:2003-05). The upper part containing the cultivation channel was manufactured by replica casting of polydimethylsiloxane (PDMS) (Sylgard 184, Dow Corning, USA) in brass replication molds. A glass cover slip (76 × 26 mm², thickness 170 µm) was bonded to the bottom of the PDMS chip via oxygen plasma treatment. The meandering channel had a rectangular cross section of 500 × 1000 µm² and a total volume of 150 µL (Figure S1 b). The straight channel for biofilm cultivation was 3 mm wide, 1 mm high and 54 mm long (Figure S1 a). The layout of the flowcell was designed in accordance with the spacing of a microwell plate, enabling optical analysis with standard microplate readers. The minimal spotsize for fluorescence measurements in the here used microplate reader (Infinite M200 pro, Tecan, Switzerland) was 3 mm corresponding to the channel width of the straight cultivation flowcell. Cannulas (Sterican, B. Braun Melsungen AG, Germany), were inserted through horizontal holes in the molds before pouring the PDMS prepolymer to serve as placeholders for the later connection channel. The PDMS was cured at 60 °C for at least 3 h. For connection to the silicone tubing (Tygon tubing R3603 (ID = 1.6 mm) Saint-Gobain, France) standard cannulas and luer lock fittings were used (Supplementary Figure S1).

### Robotic sampler

For x-y-z positioning, high precision linear stages with step motors (LIMES, Owis, Germany) were used. A dedicated motor controller (PS 90, owis, Germany) equipped with three control modules for the individual axes was used for operating the movements. The linear stages were mounted upside down on an optical portal in order to attain a robotic liquid handling arm. The probing cannula (OD = 400 µm) was mounted on a sensitive loadcell (4) (Böcker Systemelektronik, Germany) (Supplementary Figure S8). A pumping unit (5) (MicroLab 500B, Hamilton, USA) with two independent syringe modules was installed for liquid handling. The first pump module equipped with a 25 µL glass syringe (ILS, Germany) enabled precise sampling in the microliter scale whereas the second module with a 1 mL glass syringe (ILS, Germany) allowed fast flushing of the whole system. Custom made 3D printed (DesignJet Color 3D, HP, USA) accessories were organized on a 400×600 mm² metric aluminum breadboard (spacing 25 mm, Thorlabs, USA). Thereby, a customizable layout with fixed positions of all components was realized (Supplementary Figure S6). The exact positions of the fluidic chips were determined automatically by a camera. Two reference points on the chip that were directly incorporated in the replication master were recognized by the image analysis tool (AForge.NET framework, circle detection, http://www.aforgenet.com/). This allowed for the calculation of the coordinates of predefined sampling points (Supplementary Figure S7). The calibration of the camera to needle offset was done manually. The robotic sampler system was controlled by a custom-made software with a graphical user interface (Supplementary Figure S10). Further technical details of the robotic setup and the control software will be published elsewhere.

### B. subtilis and E. coli biofilms

*E. coli* DH5α and *B. subtilis* DSM1088 were cultured in the straight flowcell using LB as feed medium. Before inoculation the whole system was equilibrated in medium for at least 12 h. For the pure culture biofilms, the system was then inoculated with overnight cultures of both species, which were diluted to an optical density of 0.1 at 600 nm (OD_600_). For the two-species-biofilms, the overnight cultures were adjusted to an OD of 0.2 and pooled afterwards in a ratio of 1:1. After inoculation, the flow was halted for 1 h to allow for initial attachment of the cells. The biofilms were then cultivated at 37 °C for 12 h with a constant medium supply at 50 µL/min flowrate. In the FISH procedure, *B. subtilis* was visualized by hybridization with Alexa Fluor 488 labeled LGC354B probe35. *E. coli* was stained with Alexa Fluor 546 labeled ENT probe36.

### Enrichment of acidophilic archaea

The inoculum for the enrichment of acidophilic archaea via microfluidic culturing techniques originated from biofilms from the former pyrite mine „Drei Kronen und Ehrt” in the Harz mountains, Germany^29, 37^. In a first experiment, a section of a native biofilm, which was stored deep frozen at −80°C, was used as inoculum. The biofilm was homogenized in medium, which was used previously to enrich acidophilic archaea^38^. It was complemented with sterile filtered solutions of 90 mM ferric sulfate and 0.13 g/l yeast extract. To prevent obstruction of the microfluidic channels during inoculation, the pestled biofilm was filtered twice. The pore sizes of the filters were 70 µm and 40 µm, respectively. The biofilm-solution was inoculated under anoxic conditions using the anoxic cultivation chamber (Supplementary Figure S1h), which was flushed with an 80% N_2_/CO_2_ 20% gas mixture. The flow rate for inoculation as well as for cultivation was 10 µl/min. After 32 days the biofilm was fixed with a 4% formaldehyde solution and stained via automated CARD-FISH. The following HRP-labelled probes were used: EUB 388-I (bacteria domain, CTGCCTCCCGTAGGAGT, 20% formamide^39^) and Arch915 (Archaea domain, GTG CTC CCC CGC CAA TTC CT, 20% formamide^39^). The amplification was conducted with the dyes Alexa546 and Alexa488 and Dapi for counterstaining. Secondary cultures, highly enriched in acidophilic archaea^30^ were cultured in hungate tubes under anoxic conditions using the same medium that was used for the cultivation of the native biofilm, complemented with sterile filtered solutions of 20 mM ferric sulfate, 0.1% yeast extract and 0.1% casein. For inoculation of the chips, 6 ml of the enrichment cultures were used. The flow rate for inoculation as well as for cultivation was 2 µl/min and the incubation occurred under fully anoxic conditions with anoxic medium. After 4 months, the biofilm was fixed with 4% paraformaldehyde and stained via automated CARD-FISH as described above. For analysis, the following HRP-labelled probes were used: ARCH915 (Archaea domain, GTG CTC CCC CGC CAA TTC CT, 20% formamide^39^) and ARM980 (ARMAN, *GCC GTC GCT TCT GGT AAT, 30 % formamide*^40^). The amplification was conducted with the dyes Alexa546 and Alexa488 and DAPI for counterstaining.

### L. chromiiresistens and E. coli biofilms

Meandric chips were used to cultivate *Escherichia coli* K12 and *Leucobacter chromiiresistens*. Both organisms were pre-cultivated overnight at 30°C with LB (lysogenic broth)-medium. An inoculum suspension with an OD_600_ of 0.15 per organism was pumped through the chips for 45 min with a flow rate of 1 µl/min. Thereafter, LB medium containing 0, 1 mM, 2 mM or 3 mM potassium chromate (K_2_Cr(VI)O_4_) was pumped through the chips with a flowrate of 2.5 µl/min. Samples for 16S-sequencing were taken at day 7. Subsequently, biofilm growth was stopped by using LB-medium containing 4% formaldehyde. The biofilms were analysed by automated FISH procedure with probes Ent (5'− CCC CCW CTT TGG TCT TGC −3'^41^ conjugated with Atto488) and HGC69a (5'-TAT AGT TAC GGC CGC CGT −3'^42^, conjugated with Alexa546). All chips were counterstained with DAPI. For 16S-sequencing analysis, templates consisted of 1 µl of the robotically withdrawn samples. Primers F_NXT_Bakt_341F (5’-CCT ACG GGN GGC WGC AG-3’) and R_NXT_Bakt_805R (5’-GAC TAC HVG GGT ATC TAA TCC-3’) were used for amplification. Sequencing was conducted by IMGM Laboratories GmbH (Martinsried, Germany) on an Illumina Miseq platform, using 2×250 bp paired-end (PE) reads. Signal processing, de-multiplexing, trimming of adapter sequences was conducted by IMGM Laboratories with the MiSeq Reporter 2.5.1.3 software. Further bioinformatic analysis of the 16S rRNA amplicon sequencing (primer cutting, quality and length trimming, merging, OTU clustering, and phylogenetic analysis) was carried out with the CLC Genomic Workbench software 10.0.1 equipped with the additional microbial genomic module 2.0.

### Catalytically active E. coli biofilms

The *E. coli* BL21 (DE3) was transformed with plasmid pET_LbADH-SBP^34^ and cultivated as biofilm in the meandering flowcell in selective LB medium for 24h at 37°C. The temperature was then reduced to 30°C and the protein production was induced by addition of 0.5 mM ITPG to the cultivation medium. To evaluate the catalytic activity, the biofilm populated flowcells were placed on the deck of the robotic sampler, syringe pumps with media containing the substrate NDK (10 mM) were connected via polytetrafluorethylen (PTFE) tubing and biotransformation was carried out at 2 μL/min perfusion flowrate. After 1.5 h of equilibration, 10 μL samples were retrieved from various sampling points along the flowpath with the sampling robot. The collected samples were analyzed by chiral HPLC^33^.

## Acknowledgements

We acknowledge financial support of this work by the Helmholtz program BioInterfaces in Technology and Medicine.

## Author Contributions

S.H. developed the robotic sampler and established the microfluidic cultivation setup together with S.K. and T.K.. J.H. supervised and S.H., C.R and E.K. conceived conceptual and experimental work on the automated FISH method. T.K., S.K. and S.H. conceived the biofilm cultivation experiments. J.Ga., S.H., T.P., K.R. conducted the biotransformation study. S.H., H.H. and M.W. conceived the OCT studies. C.N., J.G., S.H, T.K., S.K. and C.R. wrote the manuscript. C.N and. J.G. planned and supervised the project. All authors discussed the results and commented on the manuscript.

